# What the Success of Brain Imaging Implies about the Neural Code

**DOI:** 10.1101/071076

**Authors:** Olivia Guest, Bradley C. Love

## Abstract

The success of fMRI places constraints on the nature of the neural code. The fact that researchers can infer similarities between neural representations, despite limitations in what fMRI measures, implies that certain neural coding schemes are more likely than others. For fMRI to be successful given its low temporal and spatial resolution, the neural code must be smooth at the sub-voxel and functional level such that similar stimuli engender similar internal representations. Through proof and simulation, we evaluate a number of reasonable coding schemes and demonstrate that only a subset are plausible given both fMRI’s successes and its limitations in measuring neural activity. Deep neural network approaches, which have been forwarded as computational accounts of the ventral stream, are consistent with the success of fMRI, though functional smoothness breaks down in the later network layers. These results have implications for the nature of neural code and ventral stream, as well as what can be successfully investigated with fMRI.

## Introduction

Neuroimaging and especially functional magnetic resonance imaging (fMRI) has come a long way since the first experiments in the early 1990s. These impressive findings are curious in light of fMRI’s limitations. The blood-oxygen-level-dependent (BOLD) response measured by fMRI is a noisy and indirect measure of neural activity (Logothetis, 2002, 2003, 2008) from which researchers try to infer neural function.

The BOLD response trails neural activity by 2 seconds, peaks at 5 to 6 seconds, and returns to baseline around 10 seconds, whereas neural activity occurs on the order of milliseconds and can be brief (Huettel, Song, & McCarthy, 2009). In terms of spatial resolution, the BOLD response may spillover millimeters away from neural activity due to contributions from venous signals (Turner, 2002). Likewise, differences in BOLD response can arise from incidental differences in the vascular properties of brain regions (Ances et al., 2009). Such sources of noise can potentially imply neural activity in regions where there should not necessarily be any.

The data acquisition process itself places limits on fMRI measurement. Motion artefacts (e.g., head movements by human subjects) and non-uniformity in the magnetic field reduce data quality. In analysis, three-dimensional images are constructed from slices acquired at slightly different times. Once collected, fMRI data are typically smoothed during analyses (Carp, 2012). All these factors place limits on what fMRI can measure.

Despite these weaknesses, fMRI has proved to be an incredibly useful tool. For example, we now know that basic cognitive processes involved in language (Binder et al., 1997) and working memory (Pessoa, Gutierrez, Bandettini, & Ungerleider, 2002) are distributed throughout the cortex. Such findings challenged notions that cognitive faculties are in a one-to-one correspondence with brain regions.

Advances in data analysis have increased what can be inferred by fMRI (De Martino et al., 2008). One of these advances is multi-voxel pattern analysis (MVPA), which decodes a pattern of neural activity in order to assess the information contained within (D. D. Cox & Savoy, 2003). Rather than computing univariate statistical contrasts, such as comparing overall BOLD activity for a region when a face or house stimulus is shown, MVPA takes voxel patterns into account.

Using MVPA, so-called ‘mind reading’ can be carried out — specific brain states can be decoded given fMRI activity (Norman, Polyn, Detre, & Haxby, 2006), revealing cortical representation and organization in impressive detail. For example, these analysis techniques paired with fMRI we can determine whether a participant is reading an ambiguous sentence, we can infer the semantic category of a word they are reading (Mitchell et al., 2004), and we can know whether a participant is being deceitful in a game (Davatzikos et al., 2005).

Representational similarity analysis (RSA), another multivariate technique, is particularly suited to examining representational structure (Kriegeskorte, Mur, & Bandettini, 2008; Kriegeskorte, 2009). We will rely on RSA later in this contribution, so we will consider this technique in some detail. RSA directly compares the similarity (e.g., by using correlation) of brain activity arising from the presentation of different stimuli. For example, the neural activities arising from viewing a robin and sparrow may be more similar to each other than between a robin and a penguin.

These pairwise neural similarities can be compared to those predicted by a particular theoretical model to determine correspondences. For example, Mack, Preston, and Love (2013) identified brain regions where the neural similarity structure corresponded to that of a cognitive model of human categorization, which was useful in inferring the function of various brain regions. The neural similarities themselves can be visualized by applying multidimensional scaling to further understand the properties of the space (Davis, Xue, Love, Preston, & Poldrack, 2014). RSA has been useful in a number of other endeavors, such as understanding the role of various brain areas in reinstating past experiences (Tompary, Duncan, & Davachi, 2016; Mack & Preston, 2016).

Given fMRI’s limitations in measuring neural activity, one might ask how it is possible for methods like RSA to be successful. The BOLD response is temporally and spatially imprecise, yet it appears that researchers can infer general properties of neural representations that link sensibly to stimulus and behavior. The neural code must have certain properties for this state of affairs to hold. What kinds of models or computations are consistent with the success of fMRI? If the brain is a computing device, it would have to be of a particular type for fMRI to be useful given its limitations in measuring neural activity.

## Smoothness and the Neural Code

For fMRI to recover neural similarity spaces, two notions of smoothness must be satisfied. Firstly, we consider *sub-voxel smoothness* and then we introduce the notion of *functional smoothness*.

### Sub-voxel Smoothness

Sub-voxel smoothness implies that neural activity varies in a smooth manner in temporal and spatial terms such that differences in activity can be measured at the level of a voxel.

Two fMRI analogues are shown in Figure 1; paralleling neurons with pixels and voxels with the squares on the superimposed grid. The left image shows a neural representation that is sub-voxel smooth. In such a spatially smooth representation, the transitions from red to yellow occur in progressive increments. Averaging within a square, i.e., a voxel, will not dramatically alter the high-level view of a smooth transition from red to yellow. Altering the grid (i.e., voxel) size will not have a dramatic impact on the results as long as the square does not become so large as to subsume most of the pixels (i.e., neurons). This result is in line with basic concepts from information theory, such as the Nyquist-Shannon sampling theorem. The key is that the red and yellow pixels/neurons are topologically organized: their relationship to each other is for all intents and purposes invariant to the granularity of the squares/voxels (for more details see: Chaimow, Yacoub, Ugurbil, & Shmuel, 2011; Freeman, Brouwer, Heeger, & Merriam, 2011; Swisher et al., 2010).

**Figure 1.**
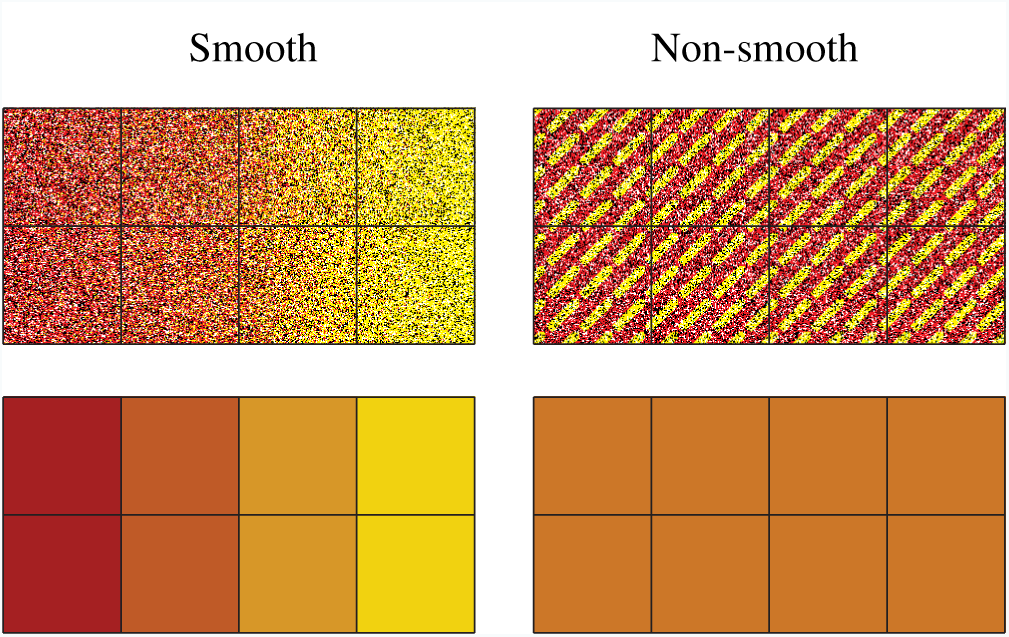
The top left quadrant depicts a spatially smooth representation at the sub-voxel level, whereas the representation to its right is non-smooth. Each square in the grid represents a voxel which averages activity within its frame as shown in the bottom panels. For the sub-voxel smooth representation, the average of each voxel (bottom left) captures the changing gradient from left to right depicted in the top-left, whereas for the non-smooth representation all voxels average to the same orange value (bottom right). Thus, differences in activation of yellow vs. red neurons would be possible to detect using fMRI for the smooth case, but not for the non-smooth case regardless of the precise boundaries of the voxels which quantize the brain.

In contrast, the right image in Figure 1 shows an inherently spatially non-smooth representation of red and yellow at the sub-voxel level. Each voxel (square in the grid), in this case, will produce an orange color when its contents are averaged. Thus, averaging the contents of a voxel in this case obliterates the representational content: red and yellow; returning instead squares/voxels that are all the same uniform color: orange. The only case in which red and yellow will be detected in a spatially non-smooth representation is when the voxel size is incredibly small, a luxury not afforded to fMRI.

Sub-voxel smoothness is consistent with proposed neural coding schemes, such as population coding (Averbeck, Latham, & Pouget, 2006; Panzeri, Macke, Gross, & Kayser, 2015; Pouget, Dayan, & Zemel, 2000) in cases where neurons with similar tunings spatially cluster (e.g., Maunsell & Van Essen, 1983). In population coding, neurons jointly contribute to represent a stimulus in much the same way as pixels were contributing to represent different colors in our toy example of a smooth representation in Figure 1. When this smoothness breaks down, similarity structures should be difficult to recover using fMRI. Indeed, a recent study with macaque monkeys which considered both single-cell and fMRI measures supports this viewpoint — stimulus aspects which were poorly spatially clustered in terms of single cell selectivity were harder to decode from the BOLD response (Dubois, de Berker, & Tsao, 2015).

The same principles extend from the spatial to the temporal domain. For fMRI to be successful, the rate of change (i.e., smoothness) of neural activity must not exceed what can be measured within a voxel. Some temporal coding schemes are not sufficiently sub-voxel smooth for fMRI to be succeed. For example, in burstiness coding, neural representations are distinguished from one another not by their average firing rate but by the variance of their activity (Fano, 1947; Katz, 1996). Under this coding scheme, more intense stimulus values are represented by burstier units, not units that fire more overall. Neural similarity is not recoverable by fMRI under a burstiness coding scheme. Because the BOLD signal roughly summates through time (Boynton, Engel, Glover, & Heeger, 1996), firing events will sum together to the same number irrespective of their burstiness.

BOLD will fail to measure other temporal coding schemes, such as neural coding schemes that rely on the precise timing of neural events, as required by accounts that posit that the synchronous firing of neurons is relevant to coding (Abeles, Bergman, Margalit, & Vaadia, 1993; Gray & Singer, 1989). Unless synchronous firing is accompanied by changes in activity that fMRI can measure, such as mean population activity, it will be invisible to fMRI (Chawla, Lumer, & Friston, 1999). As before, basic concepts in information theory, such as the Nyquist-Shannon sampling theorem, imply that temporally demanding coding schemes will be invisible to fMRI, much like how spatially non-smooth representations will be unrecoverable in fMRI (recall Figure 1).

The success of fMRI does not imply that the brain does not utilize precise timing information, but it does mean that such temporally demanding coding schemes cannot carry the day given the successes fMRI has enjoyed in understanding neural representations. Other accounts of neural firing are consistent with the success of fMRI. For example, in rate coding (discovered by Adrian & Zotterman, 1926) the frequency at which neurons fire is a function of the intensity of a stimulus. Changes in firing rate for a population of cells should be recoverable by fMRI as more blood flows to more active cells.

These examples of smooth and non-smooth representations at the sub-voxel level make clear that the neural code must be spatially and temporally smooth with respect to neural activity (which is several orders of magnitude smaller than voxels) for fMRI to be successful. Whatever is happening in the roughly one million neurons within a voxel (Huettel et al., 2009) through time is being reflected by the BOLD summation, which would not be the case if each neuron was computing something dramatically different (for in-depth discussion, see: Kriegeskorte, Cusack, & Bandettini, 2010).

### Functional Smoothness

One general conclusion is that important aspects of the neural code are spatially and temporally smooth at the sub-voxel level. In a sense, this notion of smoothness is trivial as it merely implies that changes in neural activity must be visible in the BOLD response for fMRI to be successful. As a thought experiment, an fMRI machine only capable of recording a single voxel as big as the entire brain would not be useful. In this section, we focus on a more subtle sense of smoothness that must also be satisfied, namely functional smoothness.

Neighboring voxels predominantly contain similar representations, i.e., they are topologically organized like in Figure 1 (Norman et al., 2006). However, *super-voxel smoothness* is neither necessary nor sufficient for fMRI to succeed in recovering similarity structure. Instead, a more general notion of functional smoothness must be satisfied in which similar stimuli map to similar internal representations. Although super-voxel and functional smoothness are both specified at the super-voxel level, these distinct concepts should not be confused.

To help introduce the concept of functional smoothness, we consider two coding schemes used in engineering applications, factorial and hash coding, which are both inconsistent with the success of fMRI because they do not preserve functional smoothness. In the next section, we consider coding schemes, such as deep learning networks, that are functionally smooth to varying extents and are consistent with the success of fMRI.

#### Factorial Design Coding

Factorial design is closely related to the notion of hierarchy. For example, hierarchical approaches to human object recognition (Serre & Poggio, 2010) propose that simple visual features (e.g., a horizontal or vertical line) are combined to form more complex features (e.g., a cross). From a factorial perspective, the simple features can be thought of as main effects and the complex features, which reflect the combination of simple features, as interactions.

In Table 1, a 2^3^ two-level full factorial design is shown with three factors A, B, C, three two-way interactions AB, AC, BC, and a three-way interaction ABC, as well as an intercept term. All columns in the design matrix are pairwise orthogonal.

**Table 1.** *Design matrix for a* 2^3^ *full factorial design*.

| I | A | B | C | AB | AC | BC | ABC |
| --- | --- | --- | --- | --- | --- | --- | --- |
| 1 | -1 | -1 | -1 | 1 | 1 | 1 | -1 |
| 1 | 1 | -1 | -1 | -1 | -1 | 1 | 1 |
| 1 | -1 | 1 | -1 | -1 | 1 | -1 | 1 |
| 1 | 1 | 1 | -1 | 1 | -1 | -1 | -1 |
| 1 | -1 | -1 | 1 | 1 | -1 | -1 | 1 |
| 1 | 1 | -1 | 1 | -1 | 1 | -1 | -1 |
| 1 | -1 | 1 | 1 | -1 | -1 | 1 | -1 |
| 1 | 1 | 1 | 1 | 1 | 1 | 1 | 1 |

Applying the concept of factorial design to modeling the neural code involves treating each row in Table 1 as a representation. For example, each entry in a row could correspond to the activity level of a voxel. Interestingly, if any region in the brain had such a distribution of voxels, neural similarity would be impossible to recover by fMRI. The reason for this is that every representation (i.e., row in Table 1) is orthogonal to every other row, which means neural similarity is the same for any pair of items. Thus, this coding scheme cannot uncover that low distortions are more similar to a category prototype than high distortions.

Rather than demonstrate by simulation, we can supply a simple proof to make this case using basic linear algebra. Divi<u>d</u>ing each item in the *n × n* design matrix (i.e.,Table 1) by √*n*, makes each column orthonormal, i.e., each column will represent a unit vector and be orthogonal to the other columns. This condition means that the design matrix is orthogonal. For an orthogonal matrix, *Q*, like our design matrix, the following property holds: *Q × Q*^*T*^ = *Q^T^ × Q* = *I*; where *Q*^*T*^ is the transpose of *Q* (a matrix obtained by swapping columns and rows), and *I* is the identity matrix. This property of orthogonal matrices implies that rows and columns in the factorial design matrix are interchangeable, and that both rows and columns are orthogonal.

The internal representations created using a factorial design matrix do not cluster in ways that meaningfully reflect the categorical structure of the inputs. Due to the fact that each representation is created such that it is orthogonal to every other, there can be no way for information, correlations within and between categories, to emerge. Two inputs varying in just one dimension (i.e., pixel) would have zero similarity; this is inherently not functionally smooth. If the neural code for a region was employing a technique similar to factorial design, neuroimaging studies would never recover similarity structures by looking at the patterns of active voxels in that region.

#### Hash Function Coding

Hash functions assign arbitrary unique outputs to unique inputs, which is potentially useful for any memory system be it digital or biological. However, such a coding scheme is not functionally smooth by design. Hashing inputs allows for a memory, a data store known as a hash table, that is content-addressable (Hanlon, 1966; Knott, 1975) — also a property of certain types of artificial neural network (Hopfield, 1982; Kohonen, 1987). Using a cryptographic hash function means that the arbitrary location in memory of an input is a function of the input itself.

We employed (using the same procedures as detailed below) the secure cryptographic hash algorithm 1 (SHA-1), an often-used hash function, and applied it to each value in the input vector (National Institute of Standards Technology, 2015). Two very similar inputs (e.g., members of the same category) are extremely unlikely to produce similar SHA-1 hashes. Thus, they will be stored distally to each other, and no meaningful within-category correlation will arise (i.e., functional smoothness is violated). Indeed, in cryptography applications any such similarities could be exploited to make predictions about the input.

If the neural code in a brain area was underpinned by behavior akin to that of a hash function, imaging would be unable to detect correlations with the input. This is due to the fact that hash functions are engineered in such a way as to destroy any correlations, while nonetheless allowing for the storage of the input in hash tables.

Although hash tables do not seem well-matched to the demands of cognitive systems that generalize inputs, they would prove useful in higher-level mental functions such as source memory monitoring. Indeed, to foreshadow a result below, the advanced layers of very deep artificial neural networks approximate a cryptographic hash function, which consequently makes it difficult to recover similarity structure in those layers.

## Neural Network Models

In this section, we consider whether neural networks with random weights are consistent with the success of fMRI given its limitations in measuring neural activity. Simulations in the next section revisit these issues through the lens of a deep learning model trained to classify photographs of real-world categories, such as moped, tiger, guitar, robin, etc. Each simulation is analogous to performing fMRI on the candidate neural code. These simple simulations answer whether in principle neural similarity can be recovered from fMRI data taken from certain neural coding schemes. Stimuli are presented to a model while its internal representations are scanned by a simulated fMRI machine.

The stimuli consist of simple vectors that are distortions of an underlying prototype. As noise is added to the prototype and the distortion increases, the neural similarity (measured using Pearson’s correlation coefficient ρ) between the prototype and its member should decrease. The question is whether we can recover this change in neural similarity in our simulated fMRI machine.

To help visualize the category prototype, we use a vector of pixel intensities taken from natural images (see Figure 2). This choice is simply for visualization purposes and we have obtained the same results with with categories based on Gaussian distributions (µ = 0, σ = 1) as opposed to natural images. To create input vectors from the natural images, the image intensities are mean centered (setting µ = 0) and normalized (setting σ = 1). To create a similarity structure, progressively greater levels of Gaussian noise are added to a category prototype. Twenty members per category are created by adding levels of Gaussian noise with increasing standard deviation (σ = σ_*prev*_ + 10) to each prototype (see Figure 2). Each item is re-normalized and mean centered after Gaussian noise is added, so that µ = 0 and σ = 1 regardless of the level of distortion.

**Figure 2.**
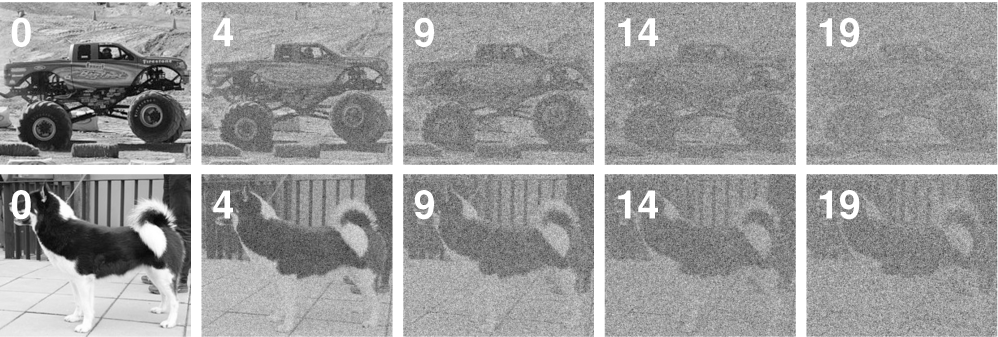
Simulations for the random neural networks involved calculating similarity between a prototype and distortions of it that were formed by adding Gaussian noise. In the figure, two prototypes are shown (labeled 0) with increasing levels of distortion shown to the right.

### Vector Space Coding

The coding schemes that follow are important components in artificial neural network models. The order of presentation is from the most basic components to complex configurations of network components. To foreshadow the results shown in Figure 3, fMRI can recover similarity structure for all of these models to varying degrees with the simpler models fairing better than the more complex models.

**Figure 3.**
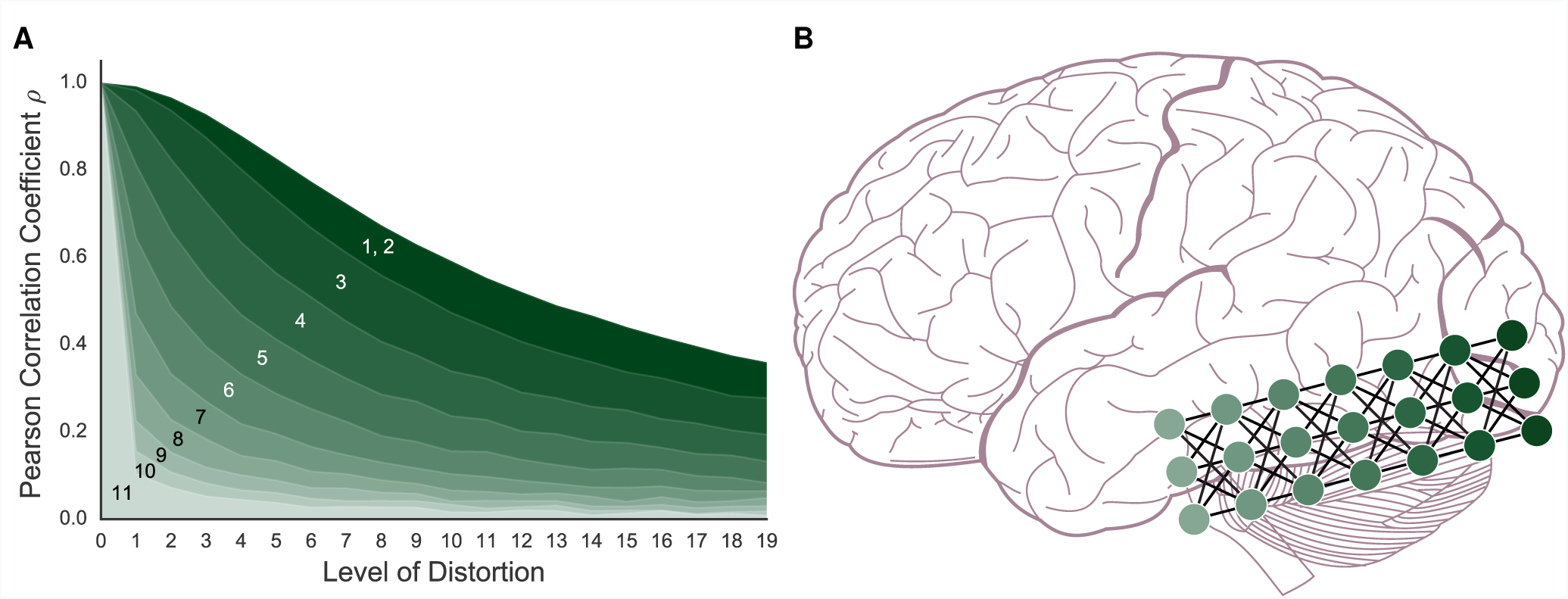
**A**: For the artificial neural network coding schemes, similarity to the prototype falls off with increasing distortion (i.e., noise). The models, numbered 1-11, are (*1*) vector space coding, (*2*) gain control coding, (*3*) matrix multiplication coding, (*4*), perceptron coding, (*5*) 2-layer network, (*6*) 3-layer network, (*7*) 4-layer network, (*8*) 5-layer network, (*9*) 6-layer network (*10*) 7-layer network, and (*11*), 8-layer network. The darker a model is, the simpler the model is and the more the model preserves similarity structure under fMRI. **B**: A deep artificial neural network and the ventral stream can be seen as performing related computations. As in our simulation results, neural similarity should be more difficult to recover in the more advanced layers.

The first in this line of models considered is vector space coding (i.e., 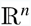), in which stimuli are represented as a vector of real-valued features (in our case pixels). Representing concepts in multidimensional spaces has a long and successful history in psychology (Shepard, 1987). For example, in a large space, lions and tigers should be closer to each other than lions and robins because they are more similar. The kinds of operations that are naturally done in vector spaces (e.g., additions and multiplications) are particularly well suited to the BOLD response. For example, the haemodynamic response to individual stimuli roughly summates across a range of conditions (Dale & Buckner, 1997) and this linearity seems to extend to representational patterns (Reddy, Kanwisher, & VanRullen, 2009).

In this neural coding scheme, each item (e.g., a dog) is represented as the set of values in its input vector (i.e., a set of numbers with range [-1, 1]). This means that for a given stimulus, the representation this model produces is identical to the input. In this sense, vector space coding is functionally smooth in a trivial sense as the function is identity. As shown in Figure 3, neural similarity gradually falls off with added distortion (i.e., noise). Therefore, this very simple coding scheme creates representational spaces that would be successfully detected by fMRI.

### Gain Control Coding

Building on the basic vector space model, this scheme encodes each input vector by passing it through a monotonic non-linear function, the hyperbolic tangent function (tanh), which is functionally smooth. This results in each vector element being transformed, or squashed, to values between [−1, 1]. Such functions are required by artificial neural networks (and perhaps the brain) for gain control (Priebe & Ferster, 2002). The practical effect of this model is to push the values in the model’s internal representation toward either 1 or 1. As can be seen in Figure 3, neural similarity is well-captured by the gain control neural coding model.

### Matrix Multiplication Coding

This model performs more sophisticated computations on the input stimuli. In line with early connectionism and Rescorla-Wagner modeling of conditioning, this model receives an input vector and performs matrix multiplication on it, i.e., computes the weighted sums of the inputs to pass on to the output layer (Knapp & Anderson, 1984; Rescorla & Wagner, 1972). These simple one-layer neural networks can be surprisingly powerful and account for a range of complex behavioral findings (Ramscar, Dye, & Klein, 2013). As we will see in later sub-sections, when a non-linearity is added (e.g., tanh), one-layer networks can be stacked on one another to build deep networks.

This neural coding scheme takes an input stimulus (e.g., a dog) and multiplies it by a weight matrix to create an internal representation, as shown in Figure 4. Interestingly, as shown in Figure 4, the internal representation of this coding scheme is completely nonsensical to the human eye and is not supervoxel smooth, yet it successfully preserves similarity structure (see Figure 3). Matrix multiplication maps similar inputs to similar internal representations. In other words, the result is not super-voxel smooth, but is functionally smooth which we conjecture is critical for fMRI to succeed.

**Figure 4.**
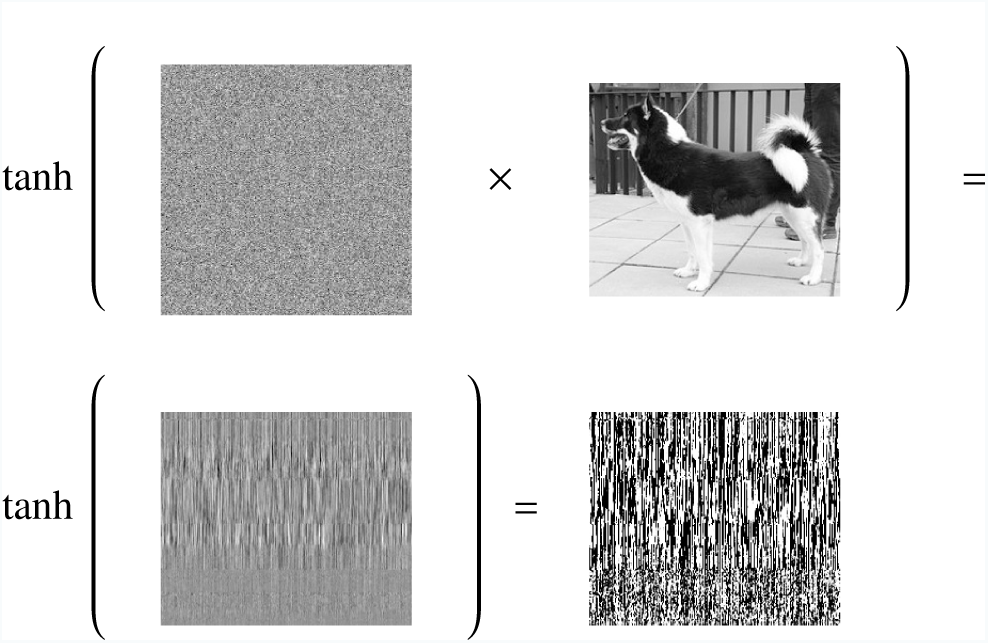
The effect of matrix multiplication followed by the tanh function on the input stimulus. The first line illustrates the stimulus being multiplied by the connection weight matrix. The output of this one-layer network is shown on the line below, as well as the outcome of applying a nonlinearity to the output of the matrix multiplication. In this example, functional smoothness is preserved whereas supervoxel smoothness is not. The result of applying this nonlinearity can serve as the input to the next layer of a multilayer network.

### Perceptron Coding

The preceding coding scheme was a single-layer neural network. To create multi-layer networks, that are potentially more powerful than an equivalent single-layer network, a non-linearity (such as tanh) must be added to each post-synaptic network layer. Here, we consider a single-layer network with the tanh non-linearity included (see Figure 4). As with matrix multiplication previously, this neural coding scheme is successful (see Figure 3) with “similar inputs lead[ing] to similar outputs” (Rummelhart, Durbin, Golden, & Chauvin, 1995, p. 31).

### Multi-layered Neural Network Coding

The basic network considered in the previous section can be combined with other networks, creating a potentially more powerful multi-layered network. These multi-layered models can be used to capture a stream of processing as is thought to occur for visual input to the ventral stream, shown in Figure 3B (DiCarlo & Cox, 2007; Riesenhuber & Poggio, 1999, 2000; Quiroga, Reddy, Kreiman, Koch, & Fried, 2005; Yamins & DiCarlo, 2016).

In this section, we evaluate whether the similarity preserving properties of single-layer networks extend to deeper, yet still untrained, networks. The simulations consider networks with 2 to 8 layers. The models operate in a fashion identical to the perceptron neural coding model considered in the previous section. The perceptrons are stacked such that the output of a layer serves as the input to the next layer. We only perform simulated fMRI on the final layer of each model. These simulations consider whether the representations that emerge in multi-layered networks are plausible given the success of fMRI in uncovering similarity spaces (see also: R. Cox, Seidenberg, & Rogers, 2015; Cowell, Huber, & Cottrell, 2009; Edelman, Grill-Spector, Kushnir, & Malach, 1998; Goldrick, 2008; Laakso & Cottrell, 2000). Such representations, as found in deep artificial neural network architectures, are uncovered by adding layers to discover increasingly more abstract commonalities between inputs (Graves, Mohamed, & Hinton, 2013; G. E. Hinton, Osindero, & Teh, 2006; G. E. Hinton, 2007; G. Hinton, Vinyals, & Dean, 2015; LeCun, Bengio, & Hinton, 2015).

As shown in Figure 3, the deeper the network the less clear the similarity structure becomes. However, even the deepest network preserves some level of similarity structure. In effect, as layers are added, functional smoothness declines such that small perturbations to the initial input result in final-layer representations that tend to lie in arbitrary corners of the representational space, as the output takes on values that are +1 or −1 due to tanh. As layers are added, the network becomes potentially more powerful, but less functionally smooth, which makes it less suitable for analysis by fMRI because the similarity space breaks down. In other words, two similar stimuli can engender near orthogonal (i.e., dissimilar) representations at the most advanced layers of these networks. In the Discussion section, we consider the theoretical significance of this result in tandem with the deep learning network results (next section).

## Deep Learning Networks

Deep learning networks (DLNs) have led to a revolution in machine learning and artificial intelligence (Krizhevsky, Sutskever, & Hinton, 2012; LeCun, Bottou, Bengio, & Haffner, 1998; Serre, Wolf, Bileschi, Riesenhuber, & Poggio, 2007; Szegedy, Vanhoucke, Ioffe, Shlens, & Wojna, 2015). DLNs outperform existing approaches on object recognition tasks by training complex multi-layer networks with millions of parameters (i.e., weights) on large databases of natural images. Recently, neuroscientists have become interested in how the computations and representations in these models relate to the ventral stream in monkeys and humans (Cadieu et al., 2014; Dubois et al., 2015; Guclu & van Gerven, 2015; Hong, Yamins, Majaj, & DiCarlo, 2016; Khaligh-Razavi & Kriegeskorte, 2014; Yamins et al., 2014; Yamins & DiCarlo, 2016).

In this contribution, one key question is whether functional smoothness breaks down at more advanced layers in DLNs as it did in the untrained random neural networks considered in the previous section. We address this question by presenting natural image stimuli (i.e., novel photographs) to a trained DLN, specifically Inception-v3 GoogLeNet (Szegedy, Vanhoucke, et al., 2015), and applying RSA to evaluate whether the similarity structure of items would be recoverable using fMRI.

### Architecture

The DLN we consider, Inception-v3 GoogLeNet, is a convolutional neural network (CNN), which is a type of DLN especially adept at classification and recognition of visual inputs. CNNs excel in computer vision, learning from huge amounts of data. For example, human-like accuracy on test sets has been achieved by: LeNet, a pioneering CNN that identifies handwritten digits (LeCun et al., 1998); HMAX, trained to detect objects, e.g., faces, in cluttered environments (Serre et al., 2007); and AlexNet, which classifies photographs into 1000 categories (Krizhevsky et al., 2012).

The high-level architecture of CNNs consist of many layers (Szegedy, Liu, et al., 2015). These are stacked on top of each other, in much the same way as the stacked multilevel perceptrons described previously. A key difference is that CNNs have more variety especially in breadth (number of units) between layers.

In many CNNs, some of the network’s layers are convolutional, which contain components that do not receive input from the whole of the previous layer, but a small subset of it (Szegedy, Liu, et al., 2015). Many convolutional components are required to process the whole of the previous layer by creating an overlapping tiling of small patches. Often (e.g., LeNet; LeCun et al., 1998), convolutional layers are interleaved with max-pooling layers, which also contain tile-like components that act as local filters over the previous layer. This type of processing and architecture is both empirically driven by what works best, as well as inspired by the visual ventral stream, specifically receptive fields (Fukushima, 1980; Hubel & Wiesel, 1959, 1968; Serre et al., 2007).

Convolutional and max-pooling layers provide a structure that is inherently hierarchical. Lower layers perform computations on small localized patches of the input, while deeper layers perform computations on increasingly larger, more global, areas of the stimuli. After such localized processing, it is typical to include layers that are fully-connected, i.e., are more classically connectionist. And finally, a layer with the required output structure, e.g., units that represent classes or a yes/no response as appropriate.

Inception-v3 GoogLeNet uses a specific arrangement of these aforementioned layers, connected both in series and in parallel (Szegedy, Vanhoucke, et al., 2015; Szegedy, Liu, et al., 2015; Szegedy, Ioffe, & Vanhoucke, 2016). In total it has 26 layers and 25 million parameters (inclusive of connection weights Szegedy, Vanhoucke, et al., 2015). The final layer is a softmax layer that is trained to activate a single unit per class. These units correspond to labels that have been applied to sets of photographs by humans, e.g., ‘space shuttle’, ‘ice cream’, ‘sock’, within the ImageNet database (Russakovsky et al., 2015).

Inception-v3 GoogLeNet has been trained on millions of human-labeled photographs from 1000 of ImageNet’s synsets (sets of photographs). The 1000-unit wide output produced by the network when presented with a photograph represents the probabilities of the input belonging to each of those classes. For example, if the network is given a photograph of a moped it may also activate the output unit that corresponds to bicycle with activation 0.03. This is interpreted as the network expressing the belief that there is a 3% probability that the appropriate label for the input is ‘bicycle’. In addition, this interpretation is useful because it allows for multiple classes to co-exist within a single input. For example, a photo with a guillotine and a wig in it will cause it to activate both corresponding output units. Thus the network is held to have learned a distribution of appropriate labels that reflect the most salient items in a scene. Inception-v3 GoogLeNet, achieves human levels of accuracy on test sets, producing the correct label in its five most probable guesses approximately 95% of the time (Szegedy, Vanhoucke, et al., 2015).

### Deep Learning Network Simulation

We consider whether functional smoothness declines as stimuli progress to the more advanced layers of Inception-v3 GoogLeNet. If so, fMRI should be less successful in brain regions that instantiate computations analogous to the more advanced layers of such networks. Unlike the previous simulations, we present novel photographs of natural categories to these networks. The key question is whether items from related categories (e.g., banjos and guitars) will be similar at various network layers. The 40 photographs (i.e., stimuli) are divided equally amongst 8 subordinate categories: banjos, guitars, mopeds, sportscars, lions, tigers, robins, and partridges, which in turn aggregate into 4 basic-level categories: musical instruments, vehicles, mammals, and birds; which in turn aggregate into 2 superordinates: animate and inanimate. We consider how similar the internal network representations are for pairs of stimuli by comparing the resulting network activity, which is analogous to comparing neural activity over voxels in RSA. Correlations for all possible pairings of the 40 stimuli were calculated for both a middle and a later network layer (see Figure 5).

**Figure 5.**
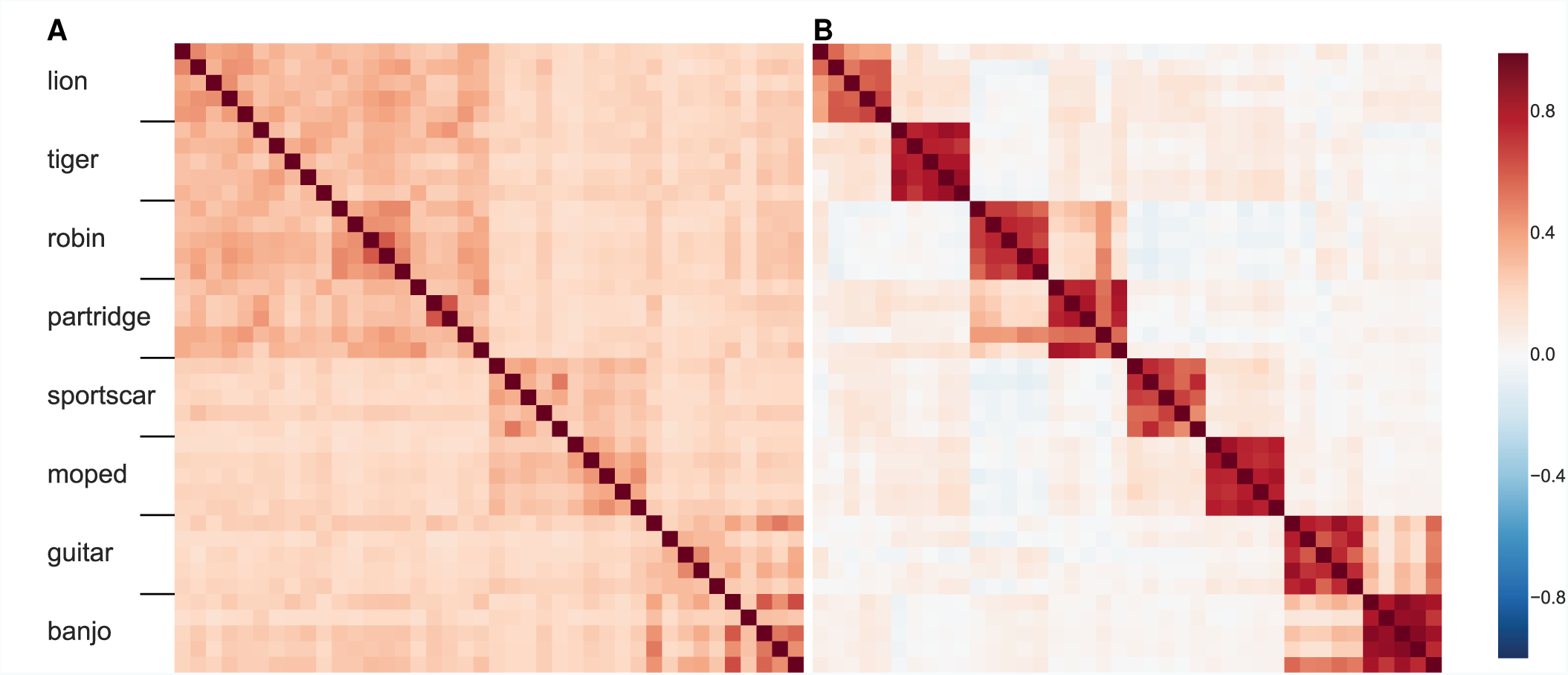
**A**: The similarity structure in a middle layer of a DLN, Inception-v3 GoogLeNet. The mammals (lions and tigers) and birds (robins and partridges) correlate forming a high-level domain. Whereas the vehicles (sportscars and mopeds) and musical instruments (guitars and banjos) form two high-level categories. **B**: In contrast, at a later layer in this network, the similarity space shows high within-category correlations and weakened correlations between categories. While some structure between categories is preserved, mopeds are no more similar to sportscars than they are to robins.

The middle layer (Figure 5A) reveals cross-category similarity at both the basic and superordinate level. For example, lions are more like robins than guitars. However, at the later layer (Figure 5B) the similarity structure has broken down such that subordinate category similarity dominates (i.e., a lion is like another lion, but not so much like a tiger). At later layers of the network (not shown), this tendency continues such that there is effectively no similarity across categories. Interestingly, the decline in functional smoothness is not a consequence of sparseness at later layers as the Gini coefficient (a measure of sparseness; Gini, 1909) is 0.947 for the earlier middle layer (Figure 5A) and 0.579 for the later advanced layer (Figure 5B), indicating that network representations are distributed in general and even more so at the later layer. Thus, the decline in functional smoothness at later layers does not appear to be a straightforward consequence of training these networks to classify stimuli, although it would be interesting to compare to unsupervised approaches of similar scale (no such network currently exists).

These DLN results are directly analogous to those with random untrained networks (see Figure 3). In those simulations, similar input patterns mapped to orthogonal (i.e., dissimilar) internal representations in later layers. Likewise, the trained DLN at later layers can only capture similarity structure within subordinate categories (e.g., a tiger is like another tiger) which the network was trained to classify. The effect of training the network was to create equivalence classes based on the training label (e.g., tiger) such that members of that category are mapped to similar network states. Violating functional smoothness, all other similarity structure is discarded such that a tiger is no more similar to a lion than to a banjo from the network’s perspective. Should brain regions operate in a similar fashion, fMRI would not be successful in recovering similarity structure therein. In the Discussion, we consider the implications of these findings on our understanding of the ventral stream and the prospects for fMRI.

## Discussion

Neuroscientists would rightly prefer a method that had both excellent spatial and temporal resolution for measuring brain activity. However, as we demonstrate in this article, the fact that fMRI has proven useful in examining neural representations, despite limitations in both its temporal and spatial resolution, says something about the nature of the neural code. One general conclusion is that the neural code must be smooth, both at the sub-voxel and functional levels.

The latter notion of smoothness is often overlooked or confused with super-voxel smoothness, but is necessary for fMRI to recover similarity spaces in the brain. Coding schemes, such as factorial and hash coding, are useful in numerous real-world applications and have an inverse function (i.e., one can go backwards from the internal representation to recover the unique stimulus input). However, these schemes are incompatible with the success of fMRI because they are not functionally smooth. For example, if the brain used such coding schemes, the neural representation of a robin would be no more similar to that of a sparrow than to that of a car. The fact that such neural similarities are recoverable by fMRI suggests that the neural code differs from these schemes in many cases.

In contrast, we found that the types of representations used and generated by artificial neural networks, including deep learning networks, are broadly compatible with the success of fMRI in assessing neural representations. These coding schemes are functionally smooth in that similar inputs tend toward similar outputs, which allows item similarity to be reflected in neural similarity (as measured by fMRI). However, we found that functional smoothness breaks down as additional network layers are added. Specifically, we have shown that multi-layer networks eventually converge to something akin to a hash function, as arbitrary locations in memory correspond to categories of inputs.

These results take on additional significance given the recent interest in deep artificial neural networks as computational accounts of the ventral stream. One emerging view is that the more advanced the layers of these models correspond to more advanced regions along the ventral stream (Cadieu et al., 2014; Dubois et al., 2015; Guclu & van Gerven, 2015; Hong et al., 2016; Khaligh-Razavi & Kriegeskorte, 2014; Yamins et al., 2014; Yamins & DiCarlo, 2016).

If this viewpoint is correct, our results indicate that neural representations should progressively become less functionally smooth and more abstract as one moves along the ventral stream (recall Figure 3). Indeed, neural representations appear to become more abstract, encoding whole concepts or categories, as a function of how far along the ventral stream they are located (Bracci & de Beeck, 2016; DiCarlo & Cox, 2007; Riesenhuber & Poggio, 1999, 2000; Yamins & DiCarlo, 2016). For example, early on in visual processing, the brain may extract so-called basic features, such as in broadly-tuned orientation columns (Hubel & Wiesel, 1959, 1968). In contrast, later on in processing, cells may selectively respond to particular individual stimulus classes (i.e., Jennifer Aniston, grandmother, concept, or gnostic cells; Gross, 2002; Konorski, 1967; Quiroga et al., 2005), irrespective of orientation, etc.

Likewise, we found that Inception-v3 GoogLeNet’s representations became symbol-like at advanced network layers such that items sharing a category label (e.g., tigers) engendered related network states, while items in other categories engendered orthogonal states (recall Figure 5). Our simulations of random networks also found reduced functional smoothness at advanced network layers, suggesting a basic geometric property of multi-layer networks. The effect of training seems limited to creating network states in which stimuli that share the same label (e.g., multiple viewpoints of Jennifer Aniston) become similar and items from all other categories (even if conceptually related) become orthogonal.

If so, areas further along the ventral stream should prove less amenable to imaging (recall Figure 3). Indeed, a recent object recognition study found that the ceiling on observable correlation values becomes lower as one moves along the ventral stream (Bracci & de Beeck, 2016).

In cognitive science, research is often divided into levels of analysis. In Marr’s levels, the top level is the problem description, the middle level captures how the problem is solved, and bottom level concerns how the solution is implemented in the brain (Marr, 1982). Given that the “how” and “where” of cognition appear to be merging, some have questioned the utility of this tripartite division (Love, 2015).

Our results suggest another inadequacy of these three levels of description, namely that the implementation level itself should be further subdivided. What is measured by fMRI is at a vastly more abstract scale than what can be measured in the brain. For example, major efforts, like the European Human Brain Project and the Machine Intelligence from Cortical Networks project (Underwood, 2016), are chiefly concerned with fine-grained aspects of the brain that are outside the reach of fMRI (Chi, 2016; Frégnac & Laurent, 2014). Likewise, models of spiking neurons (e.g., Wong & Wang, 2006) are at a level of analysis lower than where fMRI applies.

Nevertheless, fMRI has proven useful in understanding neural representations that are consequential to behavior. Perhaps this success suggests that the appropriate level for relating brain to behavior is close to what fMRI measures. This does not mean lower-level efforts do not have utility when the details are of interest. However, fMRI’s success might mean that when one is interested in the nature of computations carried out by the brain, the level of analysis where fMRI applies should be preferred. To draw an analogy, one could construct a theory of macroeconomics based on quantum physics, but it would be incredibly cumbersome and no more predictive nor explanatory than a theory that contained abstract concepts such as money and supply. Reductionism, while seductive, is not always the best path forward.

